# Exploring the impact of microbial manipulation on the early development of kelp (*Saccharina latissima*) using an ecological core microbiome framework

**DOI:** 10.1101/2023.12.13.571495

**Authors:** Jungsoo Park, Siobhan Schenk, Katherine Davis, Jennifer Clark, Laura Wegener Parfrey

**Affiliations:** Department of Botany, Biodiversity Research Centre, University of British Columbia, Vancouver, British Columbia, Canada; Department of Zoology, University of British Columbia, Vancouver, British Columbia, Canada; Cascadia Seaweed Corporation, Sidney, British Columbia, Canada

**Keywords:** microbial manipulation, core microbiome, microbial ecology, kelp cultivation

## Abstract

Kelp cultivation is a rapidly expanding economic sector, as kelp are valued for a wide range of commercial products and for restoration of kelp forest ecosystems. Microbes associated with kelp and other macroalgae play a critical role in processes such as nutrient exchange, chemical signaling, and defense against pathogens. Thus, manipulating the microbiome to enhance macroalgal growth and resilience is a promising, but largely untested, tool in sustainable kelp cultivation. The core microbiome hypothesis suggests that bacteria that are consistently found on a host (the core microbes) are likely to have a disproportionate impact on host biology, making them an attractive target for microbiome manipulation. Here, we surveyed wild *Saccharina latissima* and their surrounding environment to identify core bacterial taxa, compared to cultivated kelp, and experimentally tested the effect of cultured bacterial isolates on kelp development. We find that core bacteria are nearly absent in cultivated juvenile sporophytes in nurseries but eventually colonized after outplanting kelp to ocean farm sites. We find that bacterial inoculants can have both positive and negative effects on kelp development. In line with predictions from the core microbiome hypothesis, we find a positive correlation between the frequency of the bacterial genus in the wild and the bacterial effect on the number of sporophytes in kelp co-culture experiments.

**IMPORTANCE:** The core microbiome hypothesis suggests that symbiotic microorganisms consistently associated with hosts have functional effects on host biology and health. However, there is a lack of evidence to either support or refute this idea. This study surveys the distribution of bacteria on wild and cultivated kelp to identify the core microbiome and tests the ability of bacterial isolates cultured from the surface of wild kelp to influence kelp growth and development in laboratory microbial manipulation experiments. The frequency of bacterial genera on wild kelp was positively correlated with influence on kelp development in laboratory experiments, providing support for the core microbiome hypothesis.

## INTRODUCTION

The macroalga *Saccharina latissima* (Phaeophyceae) is one of the most important species for commercial macroalgal aquaculture in the northern hemisphere because it grows quickly and is rich in valuable organic compounds used for diverse commercial products (1–3). In Western countries, *S. latissima* has been explored as a focal species for biofuels, livestock feed, and as a component in integrated multi-trophic aquaculture (4–6). Globally, increasing consumer demand for new protein sources and climate friendly foods (7) has also been a key driver for the expansion of macroalgal cultivation outside of Asia (8, 9). However, kelp cultivation faces growing challenges. Both wild and cultivated kelp are threatened by increasing abiotic stress due to climate change (10). The intensification and global expansion of macroalgal cultivation are leading to increased disease pressure that reduce in crop yields (11–13). There is also a general need to develop strategies that enable more efficient resource use in cultivation, particularly in the nursery phage, such increasing growth rates and reducing the time to outplanting into ocean farm sites (14).

Recent studies have highlighted the importance of symbiotic bacteria associated with diverse animals, plants, and algae for host development, survival, and fitness (15–18). This recognition of the influence of microbes has prompted a wave of interest in manipulating the microbiome to alter the host in ways that benefit plants, such as increasing crop productivity or resistance to microbial pathogens in plants (19, 20). One microbiome manipulation strategy is the addition of particular microbes with beneficial effects (also called probiotics). In sustainable agriculture, addition of plant-growth promoting bacteria has been shown to reduce the need for chemical fertilizers in terrestrial agricultures (21). Similarly, addition of specific bacterial taxa can increase growth rates of microalgae (22, 23) and the macroalga *Ulva* (24), and many bacteria associated with marine algae produce the plant growth hormone Indole-3-acetic acid (IAA) (25–27). Bacterial inoculants are also being developed in finfish and shellfish aquaculture to protect against pathogens and increase growth rates (28–30), and in corals are a strategy for increasing tolerance to high temperature stress (31). In cultivated macroalgae, little work has been done on microbiome manipulation. However, in experimental lab systems, the addition of bacterial inoculants influences growth and development in the macroalgae *Ulva* sp. (32, 33), *Ectocarpus* sp. (34), and *Gracilaria* sp. (35), tolerance to low salinity stress (36), and protect against disease (37). Hence, manipulation of the microbiome in macroalgal aquaculture presents a promising but underexplored strategy (38).

Bacterial communities associated with kelp are taxonomically variable across large spatial scales (39) and across seasons (40). In kelp, early studies on *Laminaria* revealed abnormal growth in axenic conditions (41), suggesting the importance of microbes in early development. Furthermore, the addition of microbial communities from various natural seawater sources alter gametophytic size and abundance of giant kelp, *Macrocystis pyrifera* in the lab (42). These findings suggest that the kelp microbiome influences development and biology of kelp. However, there is still a limited understanding of the specific taxa within the microbiome that could be targeted for further investigation.

Harnessing the microbiome as a tool for kelp cultivation requires a better understanding of the microbiome through the kelp cultivation process. Commercial kelp cultivation starts by collecting reproductive sporophytes from the wild and releasing zoospores that settle on seedspools (twine wrapped around PVC pipe) in the nursery. Zoospores then germinate into haploid gametophytes, which produce gametes after a few weeks that cross-fertilize and develop into diploid juvenile sporophytes. Once the sporophytes are 2-10 mm in length, they are outplanted to open ocean farm sites for growth to full-size (∼2-3 meters in length) sporophytes for harvest (43, 44). The nursery stage is when growers have the most control over growing conditions, and when microbial amendments would be most feasible. Correlation between substrate coverage of *S. latissima* in nurseries and frond length in open farms (2) suggests predictive potential for later growth by microbiome manipulation. Standard protocols employ strategies to remove microbes that reduce microbial growth and biofouling, such as autoclaving, pasteurization, and seawater filtration, along with physical removal of visible fouling and wiping down with iodine solutions or weak sodium hypochlorite (44–46). However, these methods are costly, especially for small farmers (12), not all of the microbes are removed, and they do not always effectively control pathogens and fouling organisms (47). Finally, there may be overlooked consequences of removing the normal microbiome and potentially beneficial microbial symbionts. For example, some associated bacteria play a crucial role in the growth and development of a filamentous brown macroalga, *Ectocarpus* sp. (34).

The diversity of microbes associated with hosts like macroalgae and variability of the microbiome over space and time make it challenging to figure out which bacteria to target for microbial manipulations. The core microbiome hypothesis states that core bacterial taxa that are consistently associated with host organisms are more likely to influence host function (48); if supported, core taxa would be promising targets. The rationale of this hypothesis is rooted in an ecological and evolutionary perspective (49), wherein the consistent associations between hosts and bacteria may form through vertical inheritance and/or selective acquisition mechanisms. For instance, the bacterium *Roseobacter* sp., frequently found in marine algae, is attracted to the release of DMSP by the seaweed *Ulva* spp., leading to consistent associations (50, 51). *Roseobacter* sp. utilize DMSP as a food signal, consuming glycerol produced by *Ulva*, and reciprocally produces morphogens crucial for *Ulva*’s normal development (52, 53). This interaction is likely perpetuated by evolutionary adaptations, driving macroalgae to evolve mechanisms for recruiting beneficial bacteria or adapting to require bacterial byproducts, such as growth hormones and morphogens, by losing their own ability to produce these compounds. The core microbiome hypothesis is intuitive, but untested. Here, we test the core microbiome hypothesis using field surveys and laboratory experiments with *Saccharina latissima*.

In this study, we analyzed a large 16S rRNA gene dataset of *S. latissima* sampled across four sites near Vancouver, BC, Canada over time to characterize the *S. latissima* microbiome in the wild and identify core taxa as those frequently present (found on a majority of samples) and enriched compared to the environment. We then assessed the distribution of these core taxa on cultivated kelp, both in nurseries and open ocean farms. We hypothesized that the core taxa would consistently be found in cultivated kelp if they are vertically inherited or readily acquired by kelp. In Part 2, we individually co-cultured 101 isolates representing 27 bacterial genera with kelp spores and assessed their influence on kelp development. These genera are present at 0 – 94% frequency on wild sugar kelp, enabling us to test for correlations between frequency and influence on kelp development. Based on the core microbiome hypothesis, we predicted that isolates found at higher frequency would be more likely to increase host development rates.

## RESULTS

### Bacterial communities of wild and cultivated *S. latissima*

We analyzed the overall bacterial communities associated with wild and cultivated *S. latissima*, comparing them to their environment. Environmental samples allowed us to distinguish bacteria enriched on kelp from those common in the surrounding microbial pool. Samples were taken from four sites in the Salish Sea near Vancouver, BC monthly for four months (three sites) or 12 months (one site; Table 1). We found that the bacterial communities on wild and cultivated *S. latissima* differed significantly from those in neighboring environmental samples, which included rock surfaces and seawater (PERMANOVA: Pseudo-F_(2:423)_=42.631, R^2^=0.16, p<0.001) [Fig. 2A]. There were significant differences between wild and cultivated *S. latissima* at open ocean sites (PERMANOVA: Pseudo-F_(1:261)_=15.435, R^2^=0.05, p<0.001 [Fig. 2B]. Bacterial communities associated with *S. latissima* at the nursery facilities also significantly differed from wild *S. latissima* (PERMANOVA: Pseudo-F_(1:206)_=15.364, R^2^=0.06, p<0.001) [Fig. 2B]. Bacterial communities found in each of these nurseries significantly differed from one another (PERMANOVA: Pseudo-F_(2:10)_=12.742, R^2^=0.71, p<0.001).

**FIG 1.**
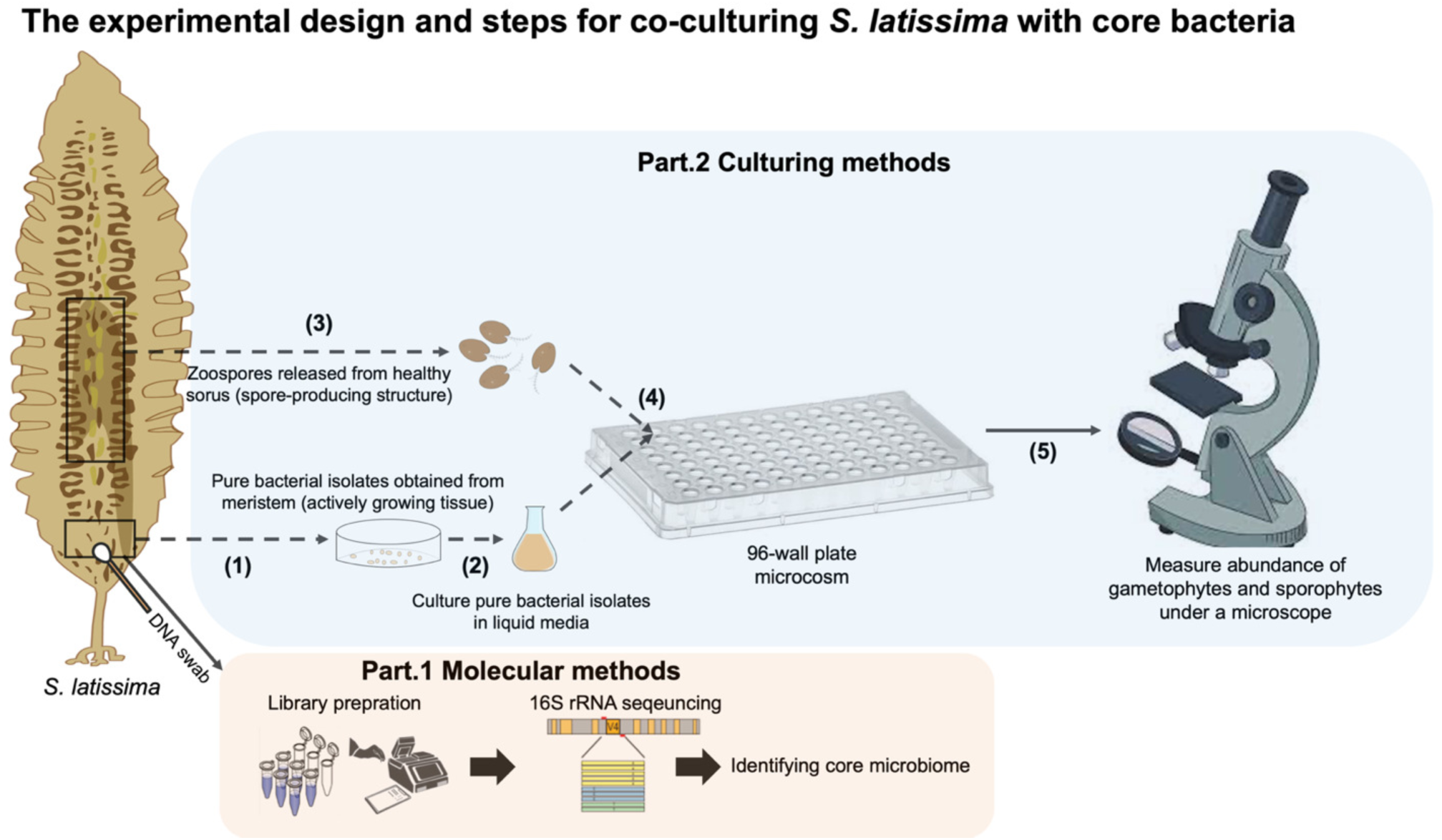
| The experimental design for co-culturing *S. latissima* with bacterial isolates. In Part 1, bacterial communities associated with *S. latissima* were analyzed using bioinformatics techniques to identify core bacterial taxa. In Part 2, purified bacterial isolates were inoculated into *S. latissima* cultures in the laboratory from the zoospore stage in the laboratory to assess the impact of different bacterial isolates on the early development of *S. latissima*.

**FIG 2.**
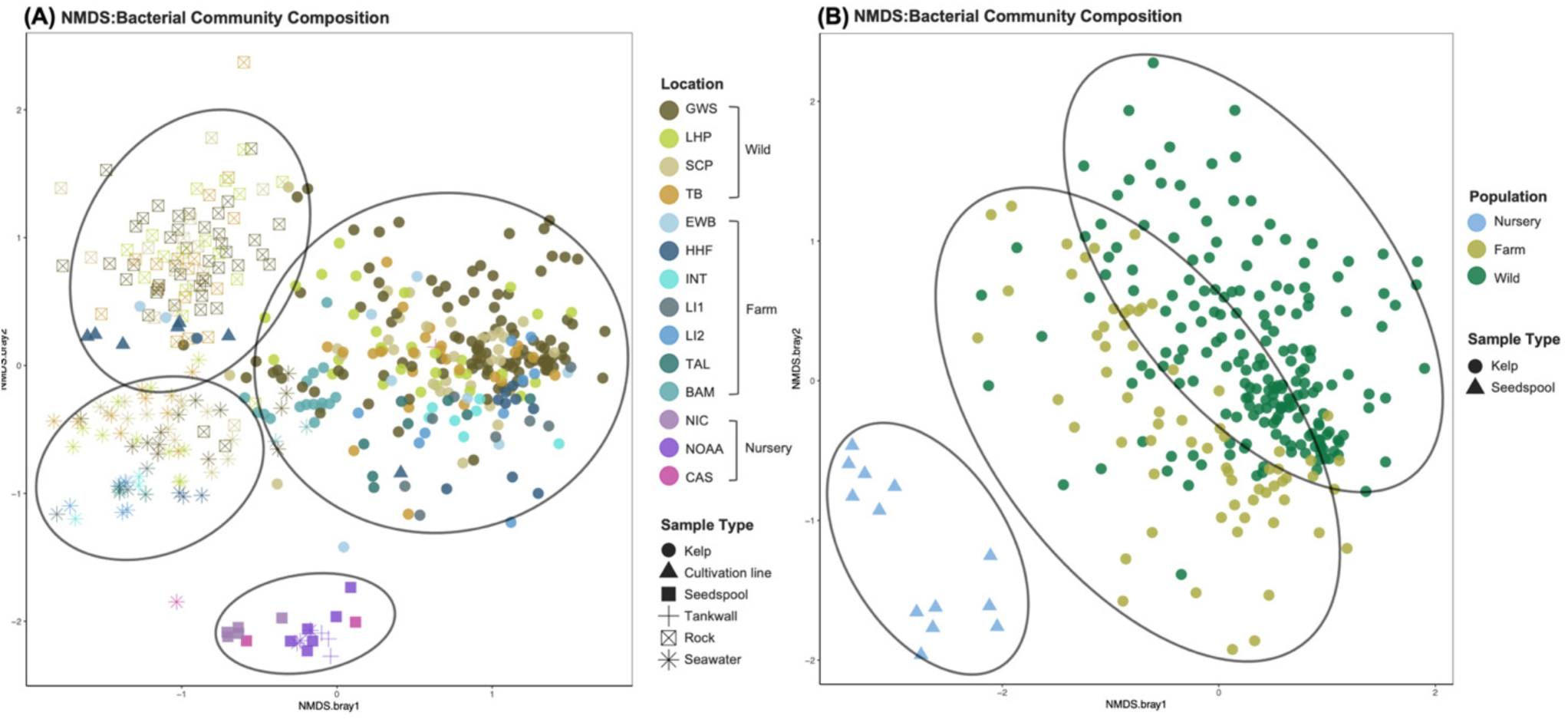
| NMDS plots were constructed using Bray-Curtis dissimilarities to compare the microbiota on. *S. latissima* **to that of the surrounding abiotic substrates.** Figure (A) shows the differences in bacterial community composition between *S. latissima* samples collected from different locations and the surrounding abiotic samples. Figure (B) illustrates the differences in bacterial community composition between *S. latissima* samples collected from different populations (wild, farm, and nursery).

**Table 1.** | Overview of samples included in the datasets. Bacterial samples included in our study sorted by sample type and location, including wild and cultivated sugar kelp (*S. latissima*) populations. The abbreviations represent sites (GWS = Girl in a Wet Suit, LHP = Lighthouse Park, SCP = Sandy Cove Park, and TB = Third Beach, BAM = Bamfield, EWB = East West Bay, HHF = Hood Head, INT = Interfor, TAL = Talbot Cove, LI = Loughboro Inlet, NOAA = WA nursery, NIC = BC nursery, Cascadia = BC nursery).

| Sample Type | Wild Site (n) |  |  |  | Ocean Farm Site (n) |  |  |  |  |  |  | Hatchery (n) |  |  |
| --- | --- | --- | --- | --- | --- | --- | --- | --- | --- | --- | --- | --- | --- | --- |
|  | GWS | LHP | SCP | TB | BAM | EWB | HHF | INT | TAL | LI1 | LI2 | NOAA | NIC | Cascadia |
| *Saccharina latissima* | 108 | 30 | 31 | 28 | 16 | 24 | 12 | 19 | 13 | 20 | 20 | 6 | 5 | 3 |
| Seawater | 14 | 11 | 13 | 10 | 2 | 3 | 5 | 3 | 4 | 3 | 4 | 3 | 0 | 1 |
| Rock | 42 | 16 | 15 | 16 | 0 | 0 | 0 | 0 | 0 | 0 | 0 | 0 | 0 | 0 |
| Cultivation line | 0 | 0 | 0 | 0 | 0 | 2 | 6 | 2 | 0 | 2 | 6 | 0 | 0 | 0 |
| Inner tank wall | 0 | 0 | 0 | 0 | 0 | 0 | 0 | 0 | 0 | 0 | 0 | 3 | 0 | 1 |

**Table 2.** | Correlation summary. Pearson’s product-moment correlation to assess the relationship between the independent factors (frequency, specificity and IndVal) of bacterial genera and the effects (log_2_ fold change) of bacterial inoculation on kelp gametophyte percent coverage and the number of sporophytes produced in each trial. Positive values of correlation indicate a positive correlation, while negative values indicate a negative correlation. Significant p-value were marked with asterisks (* or **).

|  |  |  | Frequency | Specificity | IndVal Value |
| --- | --- | --- | --- | --- | --- |
| Bacterial effects on the number of sporophytes (log2 fold change) | Trial 1 | Pearson Correlation | 0.3989687 | 0.3372198 | 0.4186339 |
|  |  | R Squared | 0.1436 | 0.1137 | 0.1753 |
|  |  | P Value | 0.00232** | 0.0976 | 0.001192** |
|  | Trial 2 | Pearson Correlation | 0.313778 | 0.3841796 | 0.3940268 |
|  |  | R Squared | 0.03835 | 0.1476 | 0.1553 |
|  |  | P Value | 0.22 | 0.1044 | 0.09507 |
|  | Trial 3 | Pearson Correlation | 0.6001401 | 0.6340727 | 0.6495264 |
|  |  | R Squared | 0.3225 | 0.402 | 0.4219 |
|  |  | P Value | 0.006596** | 0.00153** | 0.00107** |
|  | Trial 4 | Pearson Correlation | 0.7069146 | 0.486762 | 0.6185059 |
|  |  | R Squared | 0.4997 | 0.2369 | 0.3825 |
|  |  | P Value | 0.015* | 0.1289 | 0.04251* |
| Bacterial effects on the gametophyte percent coverage (log2 fold change) | Trial 1 | Pearson Correlation | 0.08397049 | 0.0666683 | 0.08344184 |
|  |  | R Squared | 0.01134 | 0.004445 | 0.006963 |
|  |  | P Value | 0.5384 | 0.6222 | 0.5372 |
|  | Trial 2 | Pearson Correlation | 0.2986401 | 0.2349035 | 0.2895252 |
|  |  | R Squared | 0.02846 | 0.05518 | 0.08382 |
|  |  | P Value | 0.2443 | 0.333 | 0.2292 |
|  | Trial 3 | Pearson Correlation | -0.2392571 | -0.1949301 | -0.2260804 |
|  |  | R Squared | 0.0017 | 0.038 | 0.05111 |
|  |  | P Value | 0.3239 | 0.3847 | 0.3117 |
|  | Trial 4 | Pearson Correlation | 0.64625232 | 0.7069834 | 0.6809983 |
|  |  | R Squared | 0.3529 | 0.4998 | 0.4638 |
|  |  | P Value | 0.03168* | 0.01498* | 0.02106* |

\* = $P \leq 0.05$ , \*\* = $P \leq 0.01$

### Core bacteria of wild *S. latissima*

We defined core bacteria of *S. latissima* as those present on the majority of wild samples The IndVal analyses of the wild *S. latissima* dataset generated a list of candidate core taxa at both the genus level (12 taxa) and ASV level (80 taxa). This analysis using a threshold of >0.7 for the IndVal index identified 8 bacterial genera and 12 bacterial ASVs, as shown in Supplementary Table 2 and 3. The core bacterial genera were *Persicirhabdus*, *Maribacter*, *Litorimonas*, *Granulosicoccus*, *Hellea*, *Cocleimonas*, *Ectothiorhodospiraceae* (genus unidentified) and *Marixanthomonas*. Additionally, there were three core ASVs belonging to *Rubritalea*, *Blastopirellula* and *Loktanella,* which were not core at the genus level. [Table. S2 and S3].

### Distribution of core bacteria: wild vs cultivated *S. latissima*

We plotted the frequency and mean relative abundance of the 12 core bacterial ASVs, as well as the sum of other ASVs within the 8 core bacterial genera, on wild *S. latissima* across four sites and over time [Fig. 3]. These four sites (GWS, TB, LHP, and SCP) are intertidal locations around Vancouver, BC, Canada, where wild *S. latissima* can be found. This allowed the visual confirmation of consistent associations with wild *S. latissima* compared to microbial communities in the surrounding environment. In most core genera, such as *Cocleimonas*, *Hellea*, and *Maribacter*, we observed a consistent presence of one predominant ASV, rather than seasonal or spatial variation in the occurrence of ASVs belonging to the same genera. This indicates that there is minimal turnover among ASVs within the same core genus across locations and over time.

**FIG 3.**
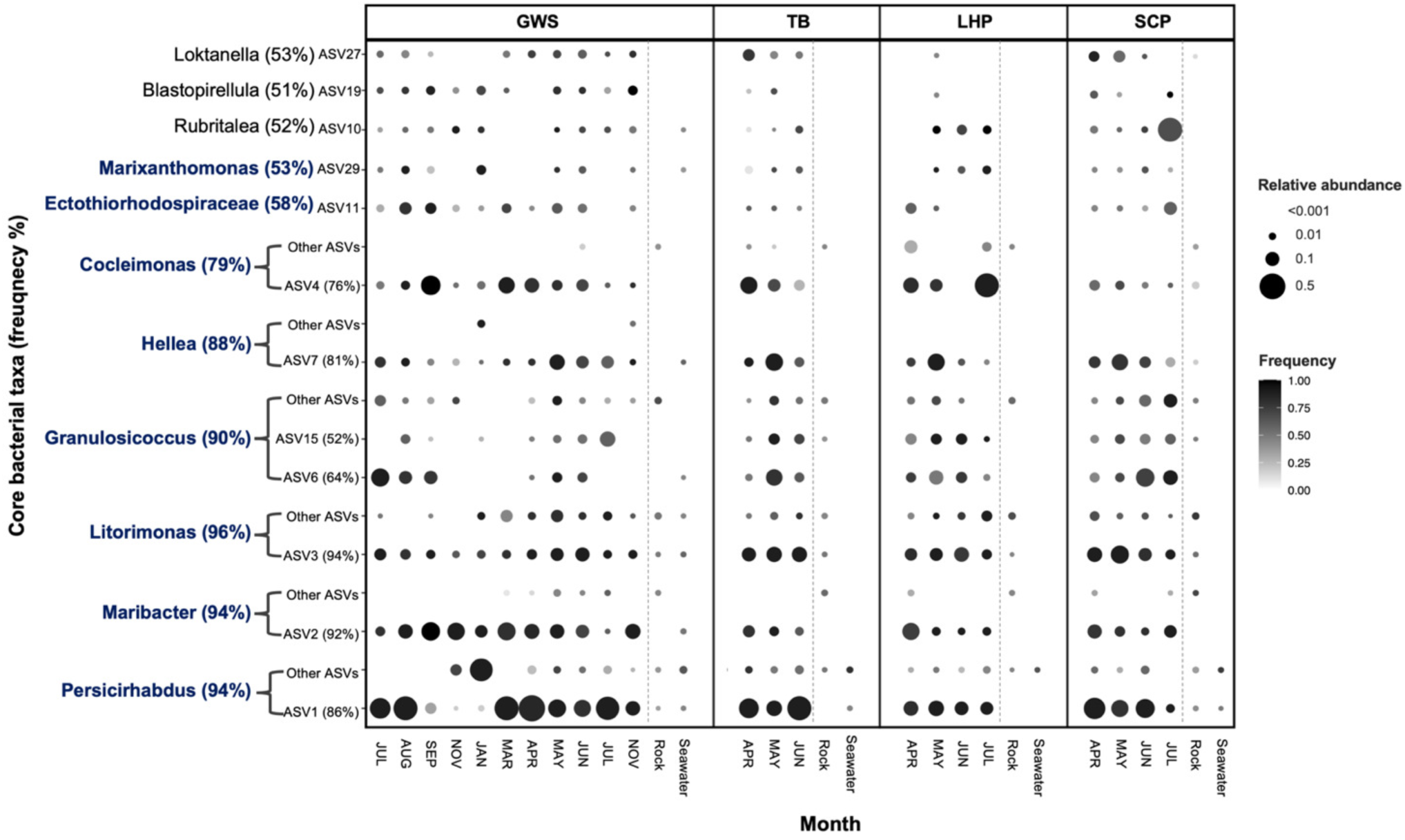
| Distribution of core taxa *S*. *latissima* in four intertidal sites (GWS, TB, LHP and SCP). The dots represent the average relative abundance and frequency of core bacterial taxa across four locations over time. The y-axis displays the core bacterial taxa identified at the genus level in blue text and at the ASV level in black text, along with the overall frequency percentage of the core bacteria. The “other ASVs” refers to summed abundance of all non-core ASVs assigned to the core bacterial genus. The monthly seawater and rock samples data were aggregated across all dates.

Furthermore, we compared those findings to the data obtained from cultivated *S. latissima* in nursery facilities and farms [Fig. 4]. These core taxa were absent on *S. latissima* seed spools at the nursery facilities, and all except *Persicirhabdus* were absent at the earliest timepoints after outplanting to some ocean sites [Fig. 4]. Interestingly, most of the core bacterial taxa colonized outplanted *S. latissima* at each of the seven open ocean farms over time. Furthermore, we observed that all core bacteria from wild kelp were found on the cultivated kelp that had been outplanted for 12 months at the Bamfield site [Fig. 4]. This finding is important because it suggests that the core bacteria identified in wild kelp populations are recruited over time.

**FIG 4.**
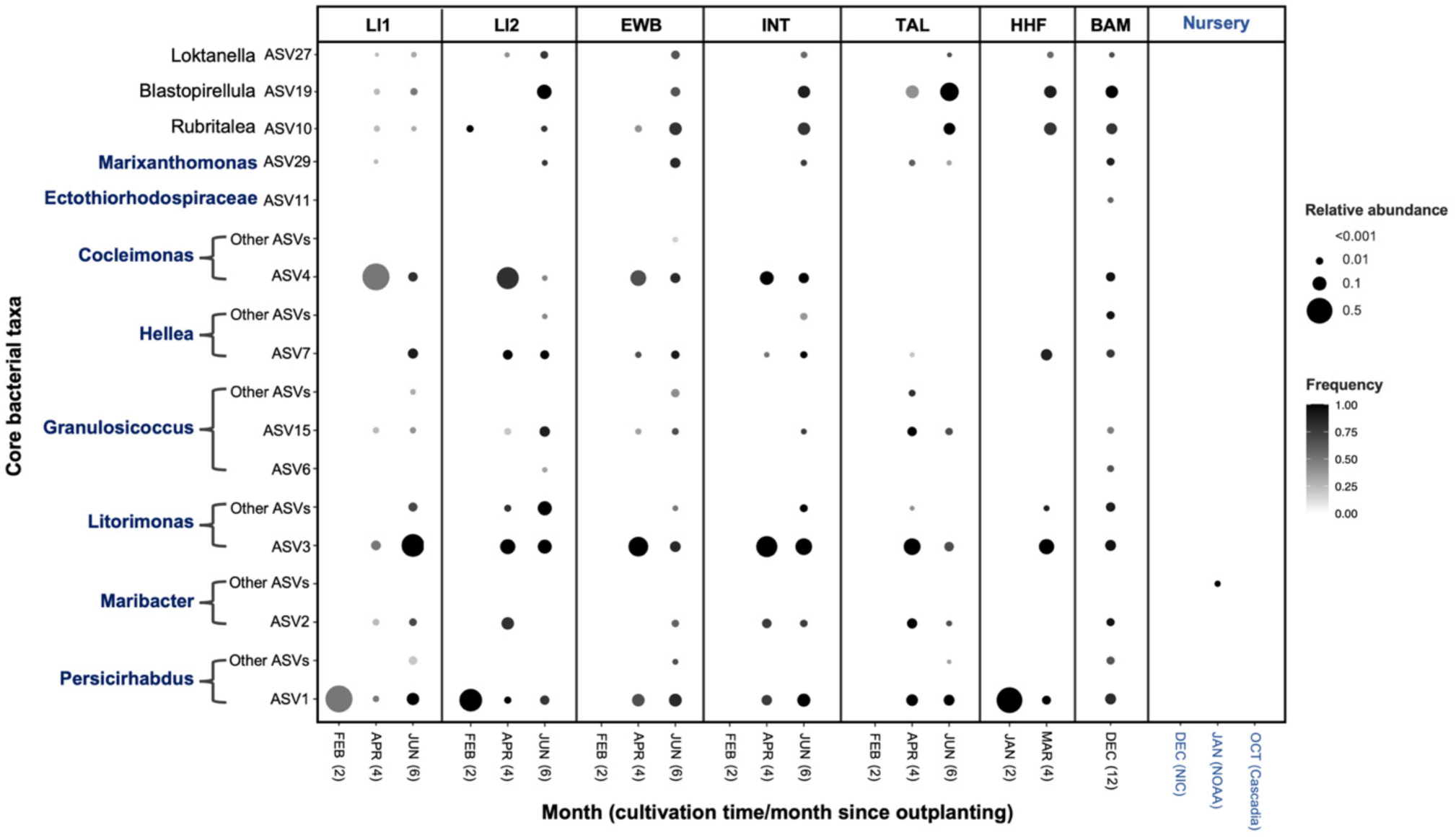
| The average relative abundance and frequency of core bacterial taxa of wild. *S. latissima* **at the genus and ASV level across seven farm sites and three nurseries over time.** The monthly associations of core bacterial taxa of wild *S. latissima* are re-evaluated in cultivated *S. latissima* at both open farms and nursery facilities. The nursery samples represent different locations and different time points. (See Fig. 3 for additional notes).

### Cultured bacterial isolates

We cultured bacteria from the surface of meristem tissue of wild *S. latissima* from the Girl in a wetsuit site. In an effort to culture as wide an array of bacteria as possible, we used a total of seven types of culture media and isolated bacteria at eight timepoints from August 2020 to February 2022 as shown in Supplementary Table 1. Isolates were identified using Sanger sequencing of the 16S rRNA gene; sequencing repeatedly failed for 27 isolates (Table S1). We obtained 127 unique bacterial isolates belonging to 32 genera. Overall, we cultured 5.9% of the bacterial genera detected in the wild *S. latissima* microbiome by 16S rRNA Illumina sequencing (32 out of 535 genera detected). Of the 32, three represented bacterial genera identified as core: *Maribacter, Hellea* and *Litorimonas*. Based on phylogenetic analyses, the isolates representing the core genera were distinct strains compared to the core ASVs. *Maribacter* isolate M16 matched ASV14, not core ASV2 (Fig. S1). *Litorimonas* isolate H37A matched ASV18 rather than core ASV3 [Fig. S1 and S2]. In addition, we found that core ASV7 within the *Hyphomonadaceae* belongs to the genus *Hellea* [Fig. 1]. We tested 101 of these bacterial isolates in co-culture experiments with *S. latissima*.

### The effects of bacterial Inoculation in *S. latissima* culture

We conducted four trials where we co-cultured *S. latissima* zoospores with bacterial isolates to observe their effect on gametophyte germination and the number of sporophytes produced. The number of isolates and replicates differed across trials (trial 1: 62 isolates, n=2 per isolate; trials 2 and 3: 25 isolates, n=4 per isolate; trial 4: 11 isolates, n=8 per isolate). Most isolates were tested only once (94 isolates), while seven were tested in multiple trials. Each experimental trial was run and analyzed separately but they are plotted together for simplicity; graphs of individual trials are in Supplementary Figure 3 and results for each isolate and trial are in Supplementary Table 4. Within each trial, the effect of bacterial isolates on kelp was calculated with t-tests comparing the log_2_ fold change of gametophyte coverage and number of sporophytes produced for each bacterial isolate to the control treatment (*S. latissima* spores without bacterial inoculants), followed by Benjamini-Hochberg correction for multiple comparisons. Across all trials, 10 isolates (9 of which were unique) had a significant positive impact on gametophyte coverage, while 3 unique isolates had a significant negative effect [Fig. 5A]. 7 isolates (6 of which were unique) had a positive impact on sporophyte number and 3 unique isolates had a significant negative effect [Fig. 5B].

**FIG 5.**
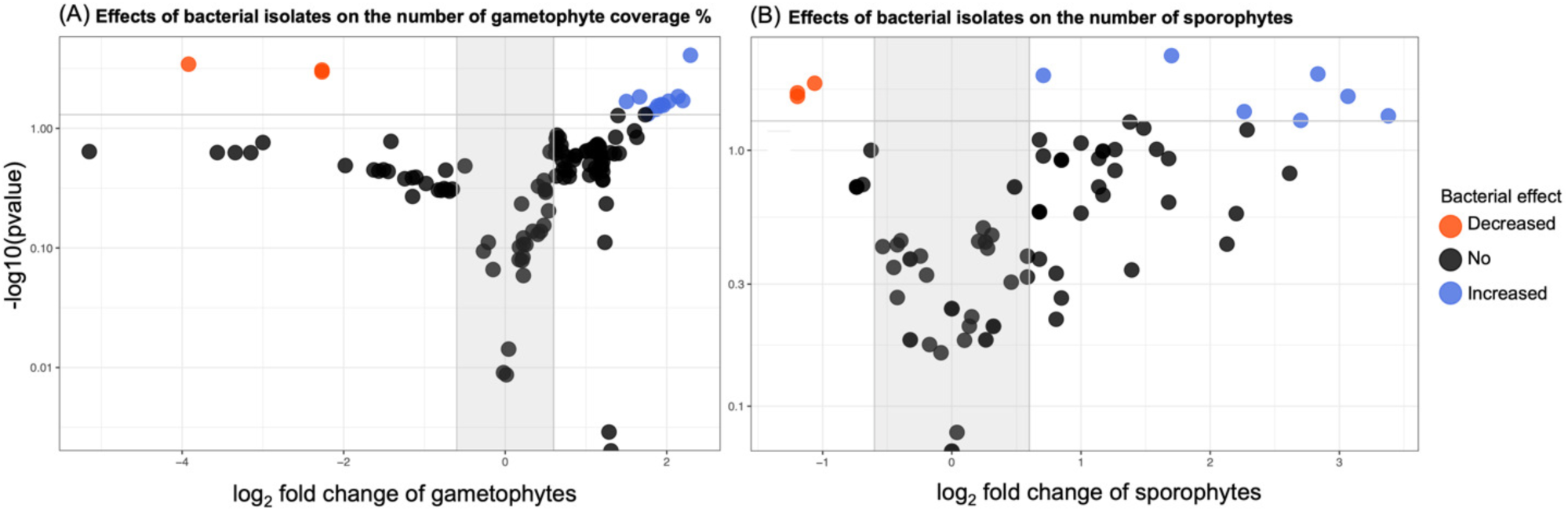
| Volcano plot depicting the effect of bacterial inoculants on *Saccharina latissima* across all four trials. The x-axis depicts the log_2_ fold change for each bacterial isolate compared to the control groups in each experimental trial on (A) gametophyte percent cover and (B) the number of sporophytes produced. The y-axis displays log_10_ corrected p-value from two-tailed t-tests comparing the control group to co-culture treatments with each bacterial isolate. Blue represents a significant positive effects, and orange represents a significant negative effect, and black represents no difference.

In trial 4 only, we assessed the persistence of the inoculated bacterium and the overall microbiome in culture wells using 16S rRNA sequencing. We confirmed that four of the seven inoculated bacterial isolates (ASV14.*Maribacter*, ASV12.*Sulfitobacter*, ASV18.*Litorimonas*, ASV5.*Pseudoalteromonas*) were indeed overrepresented over time in the kelp culture trial 4 [Fig. S5].

To assess the hypothesis that the core bacteria are more likely to influence the growth and development of kelp, we asked whether the ecological distribution of bacterial genera in the kelp microbiome correlated with their effects on *S. latissima* development in co-culture. We used the frequency, specificity, and IndVal values (specificity combined with frequency) data obtained from 16S rRNA sequencing of wild *S. latissima* in Part 1 [Table. S3] as the measures of ecological distribution and tested their relationship with co-culture results by Pearson’s product-moment correlation. These analyses were done only at the genus level because the cultured isolates largely did not correspond to ASVs detected in the 16S rRNA dataset. Frequency and IndVal value were consistently positively correlated with bacterial effects on number of sporophytes, and this correlation was significant in three of four trials [Table. 2]. Specificity showed similar trends, but the correlation was significant only in trial 3 [Table. 2]. In contrast, the relationship between ecological distribution and effect on gametophyte coverage was not consistent across trials [Table. 2]. Overall, these findings suggest that bacterial genera found at higher frequency in the kelp microbiome are more likely to induce the production of more sporophytes.

We plotted the relationship between frequency of a bacterial genus and effects of isolates from that genus in co-culture across all trials to better understand consistency across trials for a single isolate, as well as the consistency in results across different isolates from the same genus. Seven bacterial isolates were tested in three or more trials (denoted by * or + in Figure 6). Results across trial varied for these isolates but did not yield strongly contradictory results; no isolates switched from significant positive to significant negative effects or vice versa [Fig 6]. Only one isolate, *Maribacter* M16, always significantly increased the number of *S. latissima* sporophytes produced [Fig. 6B], while inoculating the remaining isolates yielded significant changes in gametophyte coverage or sporophyte numbers in some but not all trials [Fig 6]. The results for different isolates within a genus were more variable, though we note that culturing biases strongly influence the pool of cultured isolates and that trials 1 and 2 were conducted before isolates were identified, so rigorous tests are not possible. The highly culturable genera *Pseudoalteromonas*, *Shewanella*, *Sulfitobacter*, and *Vibrio* each had isolates that induced significant increases and other that induced significant decreases in the number of sporophytes (Table S4; Fig 6B). Overall, while we observed variation in effects of bacterial isolates across trials, the direction of effects generally followed consistent trends for individual isolates, but these limited data suggest more functional variability in isolates that belong to the same genus.

**FIG 6.**
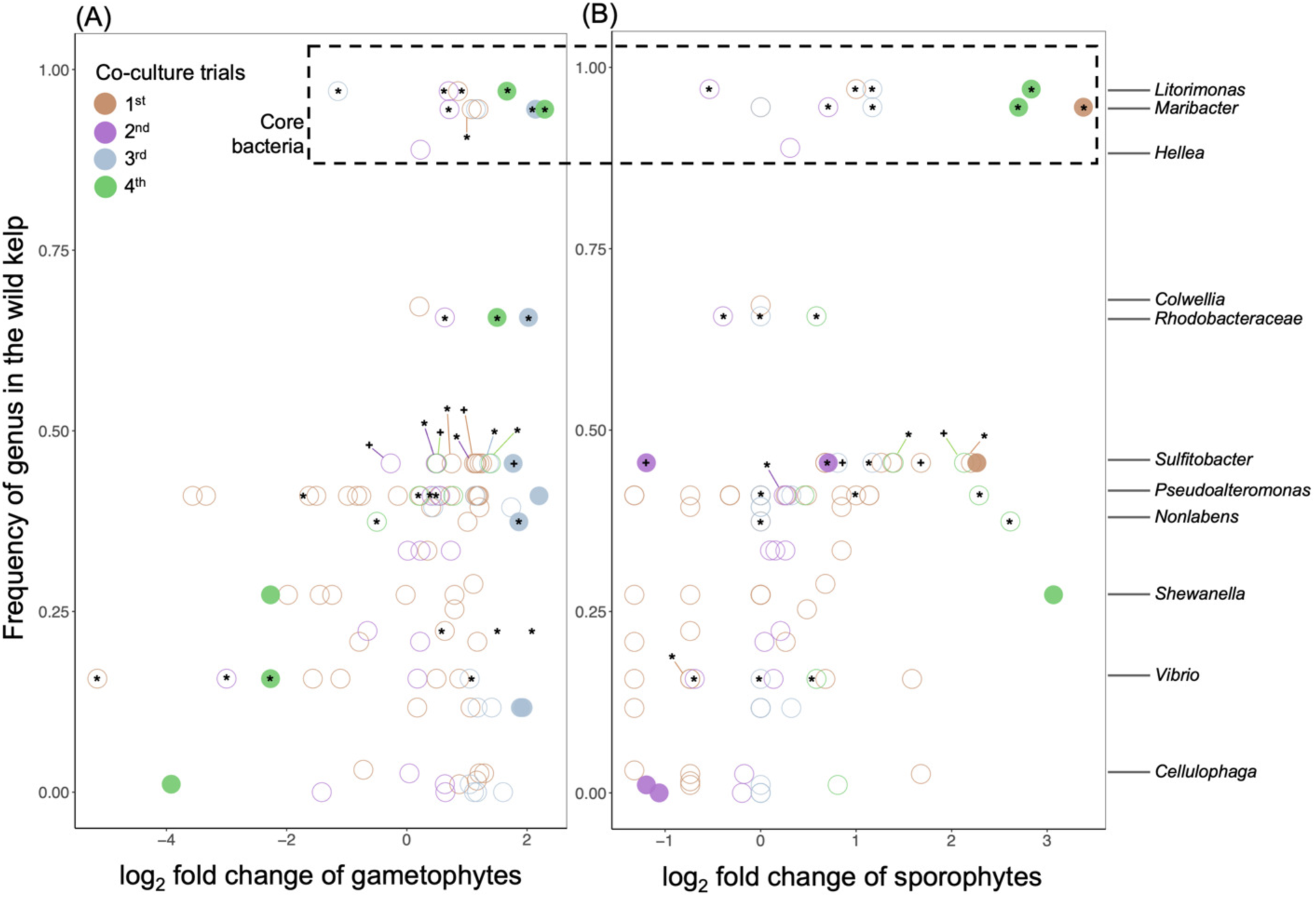
| Relationship between frequency in the wild *Saccharina* microbiome and effects of bacterial inoculation on *S. latissima* development. The x-axis displays the log_2_ fold changes for each bacterial isolate compared to the control groups in each experimental trial, for (A) gametophyte coverage % and (B) the number of sporophytes. The y-axis displays the frequency of bacterial genera identified through 16S rRNA sequencing in the wild *S*. *latissima*; example genera are labeled. Filled circles indicate significant bacterial effects (p-value < 0.05), while open circles indicate nonsignificant effects. Asterisks (*) within the circle indicate specific bacterial isolates that were tested repeatedly across four trials. Two *Sulfitobacter* isolates are indicated by asterisk (*; Mal1) and a plus sign (+; MH2).

## DISCUSSION

We investigated bacterial distribution on wild *S. latissima* and cultivated *S. latissima* and paired these data with co-culture experiments that tested the functional effect of bacteria on *S. latissima* growth and development. Together, these data enabled us to investigate the relationship between ecological distribution and functional impact, one of the first studies to empirically test this hypothesized relationship (49, 54, 55). Our findings indicate that co-cultivated bacteria can have a significant impact on the early development of *S. latissima,* and that there is a significant, positive relationship between the frequency in the *S. latissima* microbiome and the functional effect of bacterial isolates on development of *S. latissima*, in support of the core microbiome hypothesis. The high variability observed in this relationship highlights the need for further testing.

Grappling with the variability across isolates and trials is important, especially when reducing variability is also a key consideration for growers considering implementing microbial manipulation in kelp cultivation. Importantly, we did not observe significantly conflicting results (i.e. flips between significant positive and significant negative effects) across trials for the same isolate, suggesting an underlying degree of predictability. Laboratory conditions were controlled across trials, including light levels, temperature, media type, amount of starting inoculum, and duration. However, natural variation is expected across trials because *S. latissima* was collected at different time points from wild populations for each trial; *S. latissima* development may have differed as a result of seasonality, microbiome variation, host genetics, or other factors.

Earlier studies have shown seasonal variation in the chemical composition (56, 57), growth (58). The epiphytic bacterial communities also vary over time on kelp as shown in Fig. 3 and in previous research (59). Additionally, variation in the residual microbiome that was not removed by cleaning steps may also have contributed to variability across trials. While we followed protocols from industry for spore release and cultivation designed to minimize the presence of microbes (e.g., 1% betadine iodine solution on sorus tissue and autoclaving seawater media) (43, 46), these protocols do not completely remove microbes. Thus, inoculated bacteria were added to a complex community, as observed in trial 4 when the microbiome was sampled (Fig. S5). Finally, our power to detect changes in *S. latissima* development across treatments varied across trials due to differences in the number of replicates and variation in gametophyte settlement and sporophyte development across trials, as observed in variation across kelp-only controls (Supplemental table S4).

We observed several instances of contrasting effects for isolates from the same genus, including *Sulfitobacter* and *Pseudoalteromonas* [Fig. 5]. Variation across isolates is likely due to genetic differences in bacterial strains. The degree to which functional traits are phylogenetically conserved varies greatly (60), and variation in functional traits for close relatives is well known, for example in pathogenicity traits in rhizosphere *Pseudomonas* (61) and alginate metabolism in *Pseudoalteromonas* associated with giant kelp (62). One outstanding question is the relative usefulness of ecological distributions in comparison to, or paired with, trait data for expanding the catalog of bacterial inoculants and selecting local strains to be used in different regions. This will depend, in part, on the phylogenetic conservation of key traits.

Using a broadly sampled dataset of *Saccharina* from the Salish sea, we show that the microbiome of *S. latissima* both in the wild and in cultivation is distinct from the surrounding environment [Fig. 2]. We identified a small and relatively stable set of core bacteria using the definition of core taxa as those found at >50% and enriched on wild *S. latissima* compared to their surrounding environment [Fig. 3]. The overall *S. latissima* microbiome differs between cultivated and wild kelp [Fig. 2B]. The juvenile sporophytes in the nursery harbor a very different community [Fig. 2B] and lack the core taxa characteristic of the mature sporophyte blades [Fig. 4]. The core taxa appear to gradually colonize the outplanted kelp; individuals at the Bamfield location that has been outplanted for 12 months harbored all core taxa [Fig. 4]. The strikingly different community observed in the nursery phase could either be due to natural differences across development, a consequence of sterilization protocols used in cultivation. The wild kelp used to identify core taxa were from the Salish Sea near Vancouver, BC, while those used to seed nursery kelp were from Washington and the outer coast of Vancouver Island, and that nursery samples were taken in winter when prevalence of core taxa was at its lowest point [Fig. 4]. It is possible that these temporal or spatial difference shaped the nursery *S. latissima* microbiome. However, we did observe the characteristic core taxa in laboratory experimental trial 4 (with gametophytes and juvenile sporophytes), which used methods similar to those used in cultivation. It is not clear why, but core taxa present during spore release may have been transferred and better maintained in the small scale of the experiment. Further work is needed.

Our results suggest that bacterial inoculants may be useful to increase productivity in an aquaculture setting by reducing time it takes to grow kelp sporophytes to outplanting size, which is the biggest factor influencing economic viability of kelp aquaculture (14). Most bacterial inoculants were detected 28 days after inoculation (Fig. S5), suggesting that they can colonize early in development even though they are not detected in nursery samples. Further investigations are needed to understand the variability observed in response to manipulation, and to better understand the competition and colonization dynamics of inoculated bacteria. Moving beyond the early stages of kelp cultivation, it is crucial to investigate how microbial manipulation affects kelp growth and development after outplanting to open farms.

## CONCLUSION

This study demonstrates that bacterial inoculation can affect the early development of *S. latissima* and lays the foundation for developing microbial tools to enhance kelp aquaculture. The observed positive correlation between the frequency in the kelp microbiome and the effect of bacterial isolates on kelp development supports the hypothesis that core microbes are more likely to influence host function. This suggests that data on ecological distribution may be useful in the search for beneficial bacterial inoculants.

## MATERIALS AND METHODS

### Dataset description

We analyzed 16S rRNA gene amplicon datasets of the microbiome of wild *S. latissima* populations at four intertidal locations in British Columbia, Canada. We also analyzed the microbiome of cultivated *S. latissima* in three nurseries and at seven open-ocean farm sites located in British Columbia, Canada and Washington, USA [Table. 1]. We compared the kelp microbiome to samples of non-host associated microbial communities on surfaces or in the surrounding seawater to investigate bacteria that are specific to or enriched on the kelp host.

These datasets present an opportunity to identify core microbiomes of wild *S. latissima* across populations and over time, and to assess communities in the microbiome of cultivated *S. latissima* throughout the cultivation process. These datasets consist of a total of 521 samples, which contain 4,897 ASVs within 698 bacterial genera.

### Sample collection

*Wild S. latissima in coastal regions*: Microbial DNA samples were obtained from the surface of meristem tissue of *S. latissima*. [Data needed here on site lat and long, note that samples were intertidal. How many per sampling event? How frequent were sampling events? Individual kelp were rinsed with 0.22 μm filtered sterile seawater for 10 seconds to remove loosely associated microbes and then swabbed with a Puritan® sterile swab for 15 seconds. Swabs were immediately stored in 2 mL cryovials (VWR). Bare rock substrates were swabbed as a comparison to non-host associated microbial communities. Water column samples were also collected from adjacent seawater at each sampling site to characterize microbial source pool communities by filtering seawater onto a 0.22um Millipore Sterivex™ unit. All *S. latissima* tissues were promptly placed in an icebox and transported to the laboratory within two hours. Microbial samples were then collected and immediately stored at –70°C until DNA extraction.

*Cultivated S. latissima at nursery facilities*: Microbial samples were taken from three nursery facilities (NIC, NOAA, and Cascadia) at different times before they were outplanted to the open farm. These samples were collected from the nursery aquaria using the swab method, taken from the seedspool, seawater, and/or aquarium tank walls. The seedspool is an artificial substrate made up of nylon twine wrapped around a PVC pipe, on which kelp grow in commercial nurseries. The timing of collecting seedspool microbiome samples at NOAA and NIC were from juvenile sporophytes (1-4 mm blade length), just prior to outplanting into open farm ocean. Microbial swab samples from Cascadia Seaweed were obtained 3 weeks before the outplanting event which is approximately halfway through the nursery time. All microbial samples were stored at –70°C until DNA extraction.

*Cultivated S. latissima at open farm sites*: Microbiome samples were collected from outplanted kelp at seven ocean farm sites using the swab method or by extracting from whole tissue. Additionally, comparable non-host substrates (unseeded cultivation lines) were also sampled at these sites at approximately two-month intervals until harvest, except for the Bamfield site, which was only sampled once, 12 months after outplanting. All samples were stored at –70°C until DNA extraction.

### Library preparation for 16S rRNA amplicon sequencing

DNA was extracted from swabs and Sterivex filter for 16S rRNA gene amplicon sequencing using DNeasy 96 PowerSoil Pro Kit (384) (QIAGEN), following the manufacturer’s recommended protocols. PCR amplification for bacterial DNA targeted the V4 region of the 16S rRNA gene using primers, 515f: 5’– GTGYCAGCMGCCGCGGTAA–3’ and 806r: 5’ – GGACTACHVGGGTWTCTAAT –3’ (63) using dual-index barcoded primers (64). Then, we carried out amplicon library preparation, including PCR, quantification using Quant-IT Pico Green® dsDNA Assay Kit (Invitrogen). libraries were pooled at equal volumes (10 ng) of each sample, followed by DNA purification using MoBio UltaClean® PCR clean-up kit. Bioanalyzer trace was conducted at the BRC Sequencing Core, University of British Columbia to assess the quality and target size of the pooled libraries before sending the final samples to the Hakai Institute Marna Genome Lab, Quadra Island, BC for sequencing via the Illumina MiSeq Plantform (MiSeq Reagent Kit v3 (600cycle); Illumina Cat # MS-1023003).

### Bioinformatics

Demultiplexed FASTQ files were downloaded from the Illumina hub. Merging sequences from multiple datasets, quality filtering, trimming, dereplication, chimera removal, inference of ASVs and taxonomic assignment against the SILVA 138 (65) database were processed with DADA2 pipeline (66) in R environment (version 3.6.1). In the process of filtering, we discarded ASVs if they accounted for less than 0.1% of total number of reads or were found in fewer than 5 samples. Overall, we obtained 4,897 ASVs after filtering. The final products were then converted into phyloseq (version 1.28) format (67) in R for all the downstream analyses. We rarefied final 16S amplicon sequence products to 1500 reads per sample prior to beta-diversity analysis. We used nonmetric multidimensional scaling (NMDS) to visualize all samples included in this study based on Bray-Curtis dissimilarity (68). We also conducted permutational analysis of variance (PERMANOVA) using adonis2 by margin in the vegan package (version 2.5.7) (69) to test for differences among bacterial communities between *S. latissima* and environmental samples as well as between sample sites. All PERMANOVA statistics were generated with 9,999 permutations. To test the assumption of homogenous dispersion for the PERMANOVA, we conducted a beta-dispersion test with the betadipers function, also in the vegan package (data is not shown if no significant difference was found).

### Bacterial isolation

*S. latissima* tissues were collected from the “Girl in a Wetsuit” sculpture site, Stanley Park (49.302731694102754, –123.12599476839195) at 11 monthly low tide events in 2020 and 2021 and brought back to the lab for bacterial isolation within 2 hours. In the lab, meristems were rinsed with autoclaved natural seawater obtained from the Vancouver Aquarium, which has an intake pipe at 30 m depth in the Burrard inlet and is subsequently transported to and stored at The University of British Columbia. Meristems were gently rinsed with autoclaved seawater before swabbing with a Puritan® sterile swab for 15 seconds in the same manner of microbial DNA samples. Each swab was streaked on different types of customized agar plates [Table. S1] and incubated at 10°C for 4-10 days until the formation of bacterial colonies. Customization for Marne Difco Agar plate involves the incorporation of antibiotics: 1% of penicillin (100 mg/L), 1% of streptomycin (25 mg/L), and 1% of alginate acid (176.1g/mol). Additionally, we prepared custom Marine Agar Difco plates with approximately 5 g (wet weight) of ground *S. latissima* tissue per liter added to specifically target bacterial isolates unique to *S. latissima*. Lastly, we used Marine Agar Difco plates at various dilutions (1/2, 1/4, and 1/10 of the full nutrient strength) to cultivate a diverse range of bacterial isolates. Following bacterial incubation at 10°C for 4-10 days, phenotypically distinct colonies on agar plates were subcultured multiple times onto fresh standard (55.1g/L) Marine Agar Difco plates until pure cultures were obtained. All bacterial isolates obtained in this study were stored in 25% glycerol stock at –70°C and revived for cultivation when needed.

### Identification of bacterial isolates

We identified bacterial isolates using Sanger sequencing of the 16S rRNA gene. DNA was extracted from bacterial colonies on Marine Agar Difco plates using the PrepMan® Ultra Sample Preparation Reagent following the manufacturer’s standard protocols. The PCR amplification was performed with the thermocycler program: an initial DNA denaturation step at 94°C for 3 minutes, 30 cycles of DNA denaturation at 94°C for 1 minute, an annealing step at 55°C for 1 minute, an extension step at 72°C for 90 seconds, and a final extension step at 72°C for 7 minutes, using the bacterial universal primers, 515f: 5’– GTGYCAGCMGCCGCGGTAA–3’ and 1492r: 5’ – TACGGYTACCTTGTTACGACTT –3’ (70). Final PCR products were sent for Sanger Sequencing at the Genome Quebec Innovation Centre, QC, Canada.

Products were generally sequenced with both forward and reverse primers, but in some cases only forward (515F) primers yielded high quality sequence, reported in Table S1. All sequences were trimmed and quality filtered using Geneious Prime (version 2021.1). Sequences shorter than 200 base pairs were excluded. Taxonomic assignment at the genus level was carried out with the SINA aligner following standard settings (Pruesse, 2012, Bioinformatics, 28) against all databases within SINA: RDP (Ribosomal Database Project), LTP (The All-Species Living Tree Project), and SILVA (SILVA rRNA Database). Taxonomic assignment was generally The SILVA least common ancestor, except in cases where the LPT least common was more specific and LPT was used (noted in Supplemental table 1). Finally, the taxonomic assignments of isolates belonging to core genera (*Hellea, Litorimonas, Maribacter*) were refined using phylogenetic trees (see below).

### Phylogenetic analysis

To determine whether the bacterial isolates belong to the same clades as the *S. latissima* core ASVs identified from Part 1, we created phylogenetic trees for bacterial genera, using Sanger sequences from the 16S rRNA. We compared these trees to closely related sequences the SILVA 138 SSU database (65) and NCBI, which were identified using BLAST (71). We reformatted our data to FASTA files for the QIIME2 environment using the ShortRead and seqinr package in R. Sequences were aligned and the alignment masked to contain only alignment columns that are phylogenetically informative in q2-phylogeny pipeline in QIIME2 (72). Phylogenetic tree files were then constructed by using RAxML rapid bootstrap method (replicates=100) with GTRCAT model. It is important note that the resulting phylogenetic tree may not be very accurate since the sequences being compared in the analysis do not have enough informative sites to confidently distinguish between different evolutionary lineages. We also annotated the tree with information on the isolation source for each sequence using GenBank reporting of specific environment or host. Phylogenetic tree visualization and annotations were performed in Interactive tree of life (iTOL) v4 (73).

### Core bacteria of *S*. *latissima* at the genus and ASV level

To identify a suite of core bacterial taxa, we used indicator species (IndVal) (version 1.7.12) analyses using multipatt function within ‘indicspecies’ package (74) on all of the data from wild S. latissima. The IndVal metric is based on both specificity (a measure of the relative abundance of a bacterial taxon on *S. latissima* compared to environmental rock substrate and seawater samples) and frequency (a measure of its presence across *S. latissima* samples) of bacterial taxa (75, 76). Permutation tests were used to evaluate the statistical significance of the association between bacterial taxa and wild *S. latissima*. Our previous study comparing methods for identifying the core (77) led us to adopt this approach because we observed that the most consistent taxa were captured by this approach. We defined the *S. latissima*-core bacterial taxa based on an IndVal value of a 0.7 or above, with a frequency of greater than 50%, meaning the taxon was present in more than half of the macroalgal samples. Separate IndVal analyses were run to identify potential core taxa at the genus and ASV level. We then analyzed the relative abundance and frequency of the core taxa identified in wild *S. latissima* populations in the 16S amplicon sequencing data collected from cultivated *S. latissima*. We visualized the distribution of the core bacteria across sampling sites over time in both wild and cultivated *S. latissima* using the ggplot2 package (version 3.3.6) (78) in R.

### Co-cultivation for *Saccharina latissima* and isolated bacteria

*Cultivation of S. latissima:* We conducted four trials co-cultivating *S. latissima* with single bacterial isolates in Nov 2020, Jan 2021, April 2021, and Oct 2021. Each well of the 96 well plates were filled with kelp zoospores (preparation described below), a single bacterial isolate (preparation described below), and F/2 media to a final volume of 200 μL. Plates were incubated at 10°C under cool-white LED lamps (75 μmol photons m^-2^ s^-1^) with a 16h:8h light:dark cycle for 28 days. Water changes occur on day 14 during cultivation by carefully inverting the 96-well plate and refilling each well with fresh F/2 media.

For each of the four trials, we collected *S. latissima* blades with mature sorus patches from intertidal populations at the “Girl in a Wetsuit” site at low tide and transported them back to the lab for spore release within 2 hours in a cooler filled with unfiltered seawater collected at the site. We released spores according to published methods (44). Briefly, we excised mature sorus patches from the kelp blade with a sterile razor blade and rinsed the sorus tissue with autoclaved seawater. We then disinfected the sorus tissue with a 2% betadine solution for one minute, rinsed it with autoclaved seawater, and wrapped it in paper towels to dry overnight in the dark at 10°C. The next day, we submerged the dehydrated sorus tissue in autoclaved seawater for approximately 15 minutes or until the seawater turned brown and cloudy to induce spore release by osmotic shock. We counted the motile zoospores over a hemocytometer to calculate their concentration, and then plated approximately equal densities (∼5 motile zoospores/10 μL final volume) into each well of a 96-well culture plate; this was typically less than 5 μL zoospore solution and 195 μL F/2 media. We note that non-motile zoospores were also transferred to the 96-well plate and may have increased the viable cell counts.

To prepare the bacterial inoculum of each isolate we transferred a fresh colony from agar plates to liquid marine broth media (Difco) and incubated it without shaking in 13 mL screw cap tubes at room temperature for 72 hours. We then harvested the bacterial cells by centrifugation (3700 × g at 4°C for 10 min) and washed them three times with autoclaved F/2 medium, then added 3mL of sterile F/2 medium to the pellet. We then used spectrophotometry to estimate the concentration of bacterial cells by measuring their optical density at 600 nm (79). We then added a single bacterial inoculum to each of the culture wells, resulting in a final bacterial density of ∼1 × 10^7^ cells mL^-1^. The control group for each trial consisted of zoospores and F/2 medium without any added bacterial cells. The number of isolates and replicates vary across trials as follows: 62 isolates (n = 2) in trial 1, 25 isolates (n = 4) in trial 2, 25 isolates (n = 4) in trial 3, and 11 isolates (n = 8) in trial 4.

The objective of the exploratory co-culture experiments was to identify bacterial isolates that influence the biology of kelp. A total of 127 bacterial isolates were obtained based on phenotypical differences in colony appearance. Out of these, 101 bacterial isolates were tested in four co-cultivation trials with kelp [Table. S4]. We retested seven bacterial isolates that had large positive or negative effect on gametophyte coverage or the number of sporophytes during the initial co-culture trials, in order to assess the reproducibility of their effects: M16.*Maribacter*, Mal1*.Sulfitobacter*, MH2*.Sulfitobacter*, Man4.*Pseudoalteromonas* and 2Man3.*Vibrio* in trials 1-4, and 37A.*Litorimonas* and KM3. *Rhodobacteraceae* in trials 2-4.

### The impact of bacteria on *S. latissima*

To investigate the effect of bacterial isolates on the early life stages of *S. latissima*, we estimated the percent coverage of gametophytes and counted the number of sporophytes that developed. After 28 days of cultivation, we examined the entire 96-well plate under the inverted microscope. We did not systematically record the timing of gametophyte germination or sporophyte appearance. However, we observed that germination to gametophytes typically occurred after 1-2 weeks of culturing and sporophyte development took place after 3-4 weeks. These times are comparable to those reported for standard commercial nursery practices (43).

Previous studies have indicated that in open farm conditions, the coverage of both sporophytes and gametophytes on seedstring twine during the nursery phase is correlated with the frond length of *S. latissima* at harvest (2), suggesting our during the early life stages may be useful in predicting harvest outcomes.

To evaluate the differential effects of bacteria on *S. latissima*, we calculated the log_2_ fold-change for each well compared to the trial average for *S. latissima*-only controls for both the percent coverage of gametophytes and the number of sporophytes. Log_2_ fold-change cannot be calculated for values of 0 as was occasionally the case for sporophyte numbers, thus we added 1 to the number of sporophytes prior to comparison. We then used two-tailed t-tests followed by Benjamini-Hochberg correction for multiple comparisons to assess the differences between each bacterial isolate treatment and the S. latissima-only control group.

In trial 4 only, used 16S rRNA gene amplicon sequencing was used to assess the bacterial community during the experiment. This allowed us to assess whether the bacterial cultures added on day 1 persisted over time. We sampled the microbiome of culture wells by swabbing the entire well destructively (culture media, well surfaces, and *S. latissima* in culture) at day 1, day 7, day 14, day 21, and day 28 for eight treatments, including the control.

### Data Availability

Raw sequence files and metadata files have been deposited in the European Nucleotide Archive under the project accession PRJEB64485. All relevant raw data for bacterial isolates are within the main paper and its Additional supplementary tables 1, 2, 3 and 4.

## ACKNOWLEDGMENTS

We respectfully acknowledge that this research was conducted on the traditional and unceded territory of the Coast Salish Peoples, including the Squamish (Skwxwú7mesh), Musqueam (xʷməθkʷəy̓əm), and Tsleil-Waututh (səl̓ilwətaɁɬ) Nations, upon whose land Stanley Park is situated.

J.P and L.W.P designed the study. J.P and S.S collected the wild kelp microbiome data at the GW site in Stanley Park, BC, Canada. S.S collected the wild kelp microbiome from LHP, TB, and SCP. K.D and S.S also provided cultivated kelp microbiome data from the nursery and aquaculture, with support from J.C, a collaborator from Cascadia Seaweed. J.P conducted the molecular laboratory work for the partial wild kelp microbiome data and carried out all bioinformatics analyses using the combined datasets. J.P isolated and identified all bacterial cultures used in the laboratory experiment. With guidance from L.W.P, J.P wrote the manuscript and created the figures.

---

**FIG S1.**
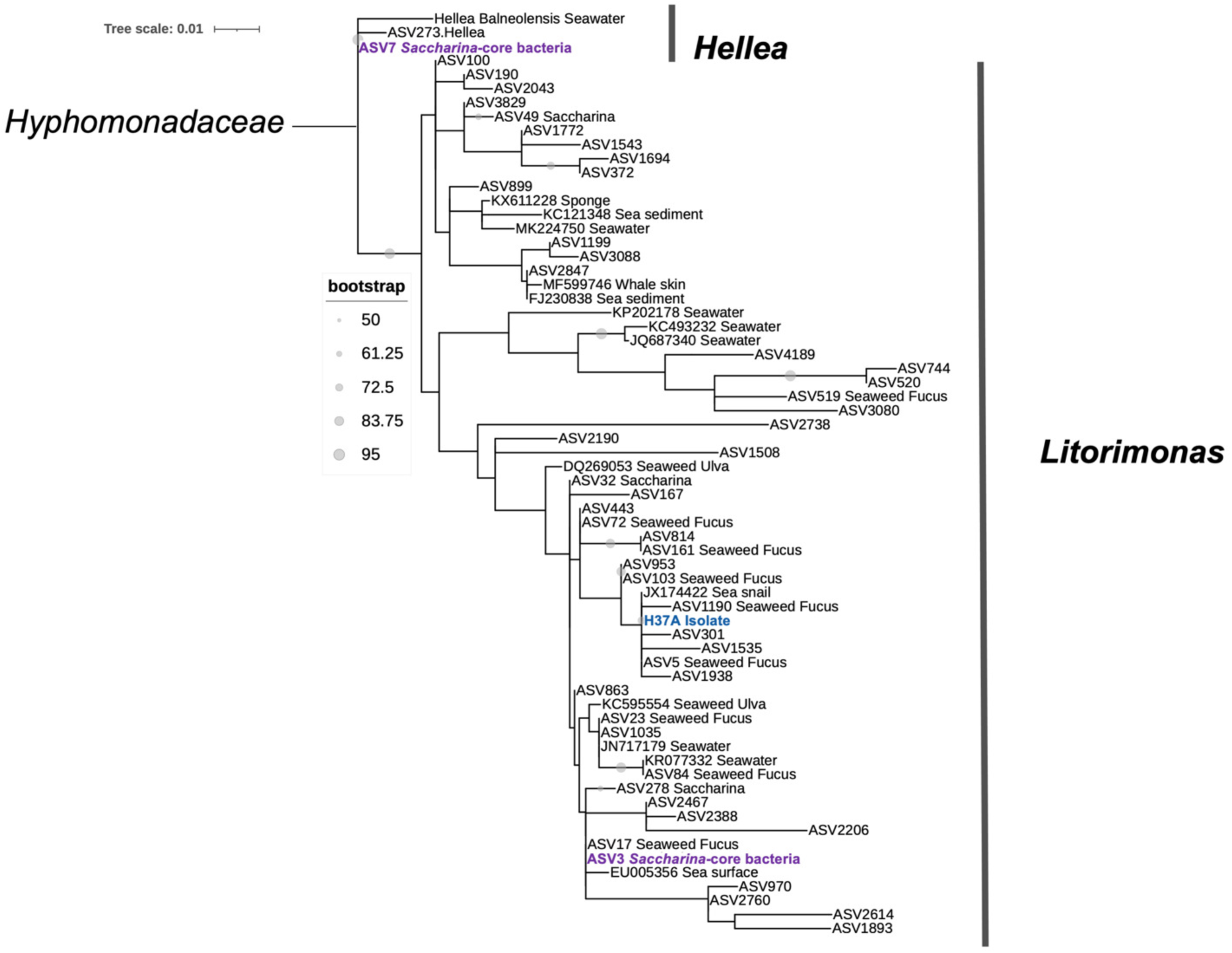
| A phylogenetic tree of *Hyphomonadaceae* clade bacterial sequences. The tree was constructed using RAxML and rapid bootstrapping. The core ASVs of *S. latissima*, identified in this study are colored purple, while the sequences of the cultured bacterial isolates in the laboratory are colored blue. Other sequences obtained from GenBank are annotated with information on the isolate source. Only bacterial sequences within branches with bootstrap values greater than 50 are shown.

**FIG S2.**
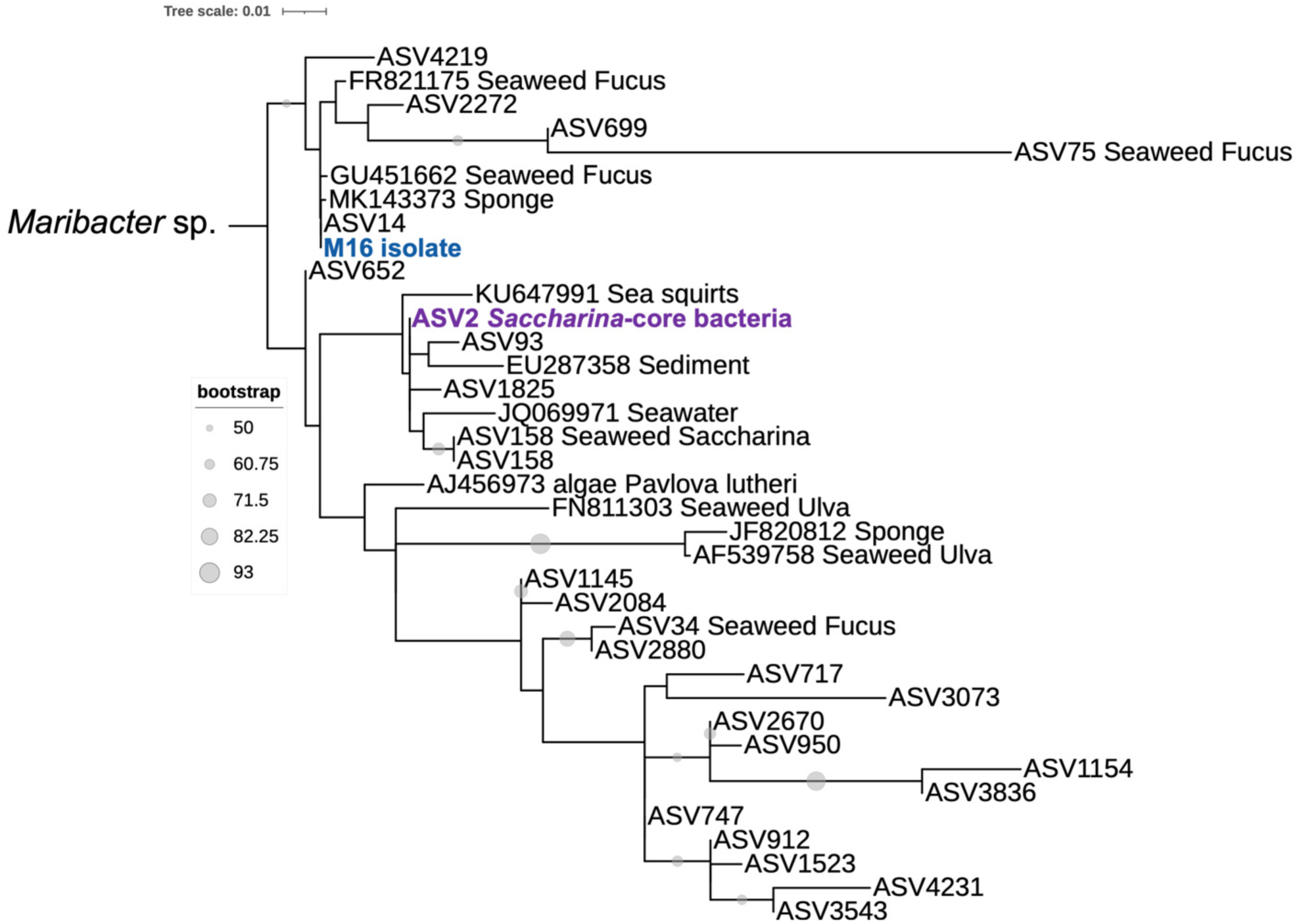
| A phylogenetic tree of *Maribacter* clade bacterial sequences. Other notes as in Fig. S1.

**FIG S3.**
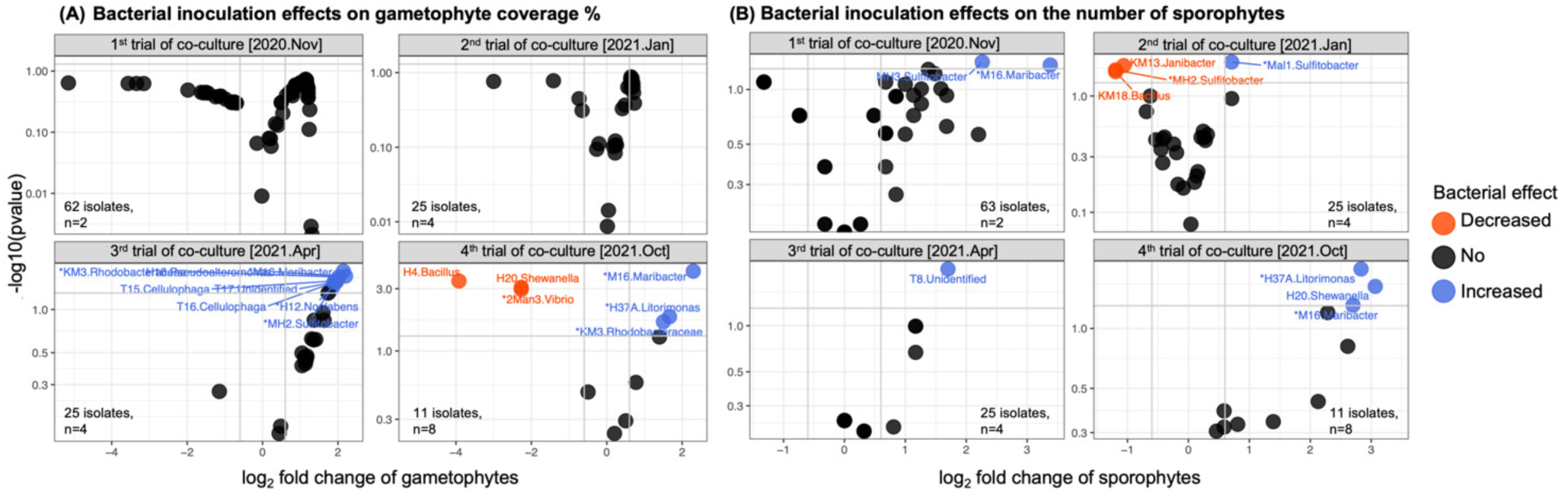
| The dot plots illustrate the effects of bacterial inoculation on *Saccharina latissima* for each individual trial. Bacterial isolates have a significant positive or negative effect on gametophyte coverage % and the number of sporophytes. Points are labelled by the strain identifier and genus of the inoculum. Other notes are in Fig. 5.

**FIG S4.**
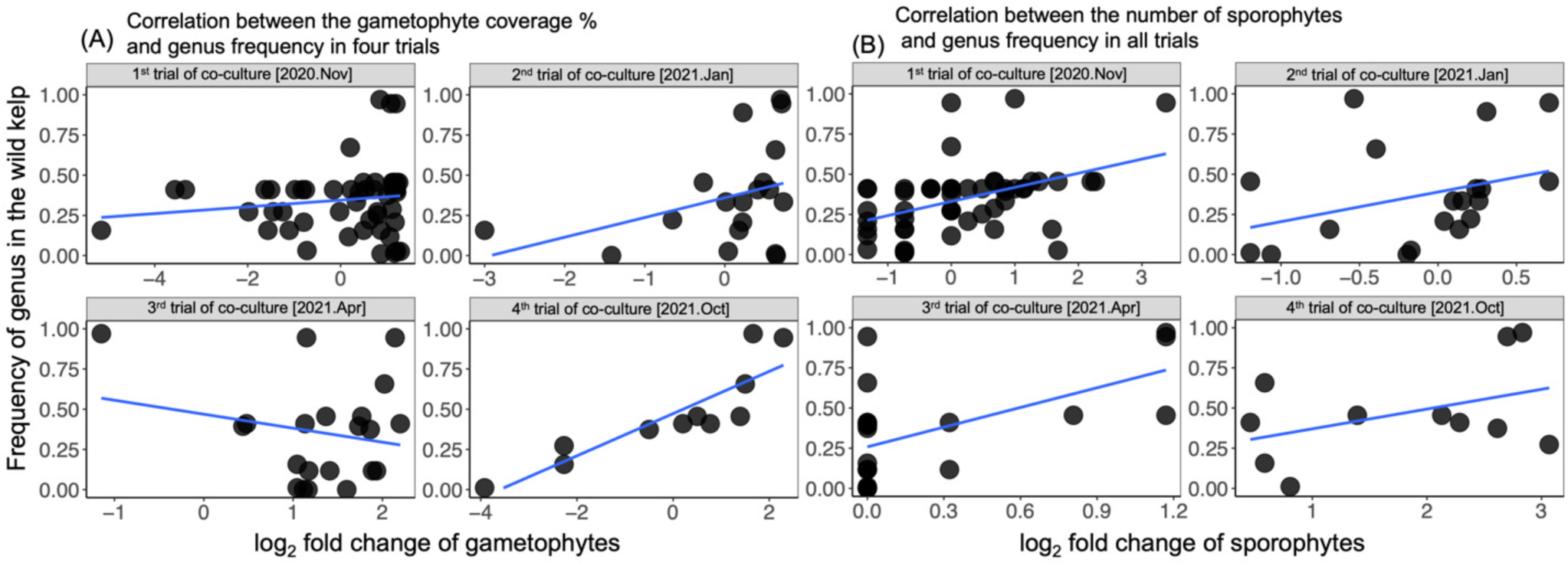
| Relationship between genus frequency on wild *S. latissima* and the effects of co-culturing bacterial isolates on *S. latissima* in the lab. The x-axis displays the log_2_ fold changes for each bacterial isolate compared to the control groups in each experimental trial, on (A) gametophyte coverage % and (B) the number of sporophytes. The y-axis displays the frequency of bacterial genera on the wild *Saccharina* identified by Illumina sequencing. Blue trend lines indicate Pearson9s product-moment correlation.

**FIG S5.**
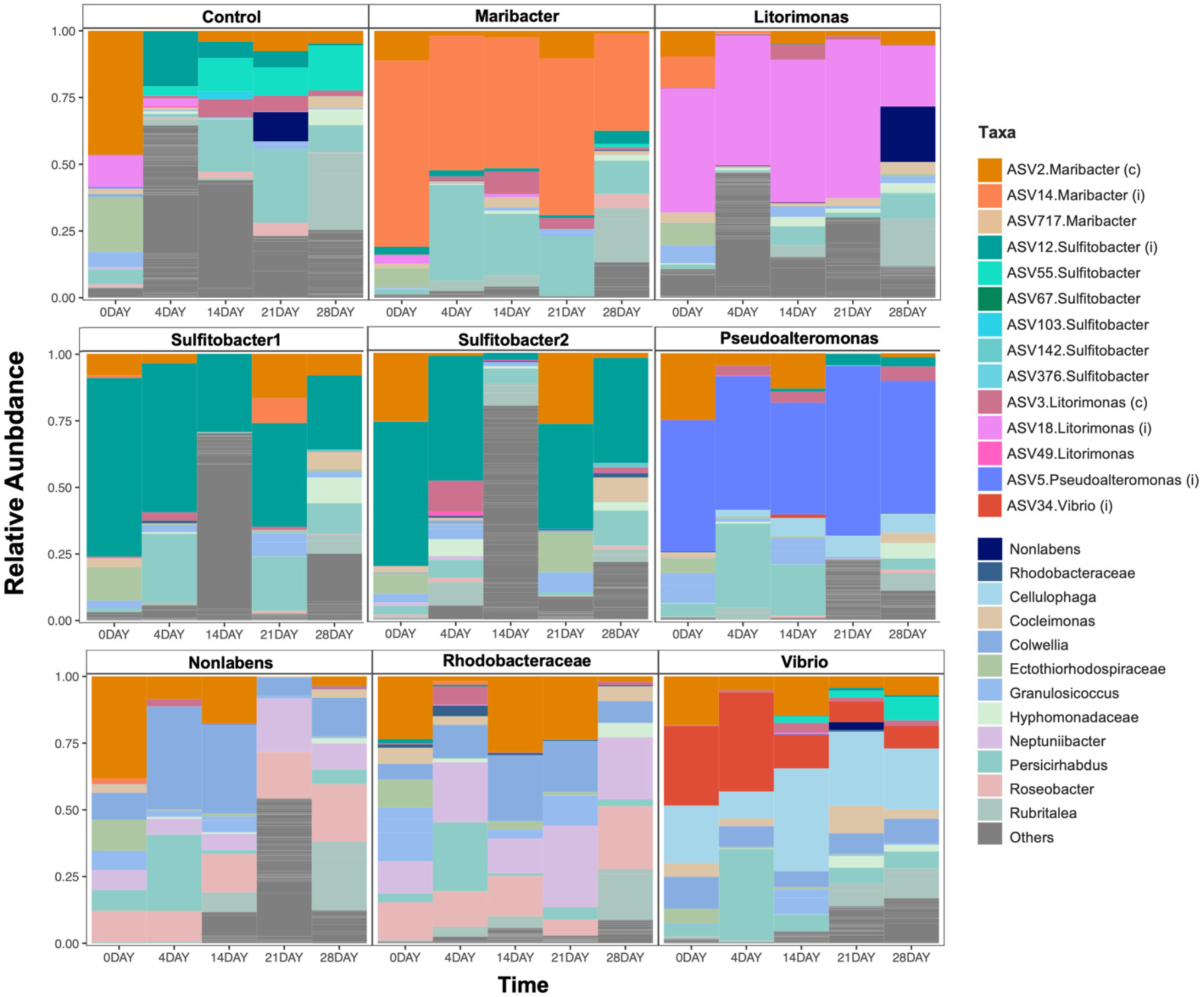
| Taxonomic bar plot of bacterial community in co-culture trial 4 over the 28-days. Most abundant genera across all trials plus Data from amplicon sequencing. The bacterial isolated that were inoculated are indicated (i) used for microbial manipulation are displayed at the ASV level to differentiate between different species within the core genera. The core ASVs (c) identified by IndVal analysis that were associated with wild S. latissima were also present in this laboratory-based microcosm. A water change occurred prior to the 14-day sampling.

